# Chromosome 8 gain drives cancer progression by hijacking the translation factor 4E-BP1 sensitizing for targeted CDK4/6 inhibition

**DOI:** 10.1101/2022.12.11.519935

**Authors:** Cornelius M. Funk, Anna C. Ehlers, Martin F. Orth, Karim Aljakouch, Jing Li, Tilman L. B. Hölting, Rainer Will, A. Katharina Ceranski, Franziska Willis, Endrit Vinca, Shunya Ohmura, Roland Imle, Jana Siebenlist, Angelina Yershova, Maximilian M. L. Knott, Felina Zahnow, Ana Sastre, Javier Alonso, Felix Sahm, Heike Peterziel, Anna Loboda, Martin Schneider, Ana Banito, Gabriel Leprivier, Wolfgang Hartmann, Uta Dirksen, Olaf Witt, Ina Oehme, Stefan M. Pfister, Laura Romero-Pérez, Jeroen Krijgsveld, Florencia Cidre-Aranaz, Thomas G. P. Grünewald, Julian Musa

## Abstract

Chromosome 8 (chr8) gains are common in cancer. However, their potential contribution to tumor heterogeneity is largely unexplored. Ewing sarcoma (EwS) is characterized by pathognomonic *FET::ETS* fusions but a general paucity of other recurrent somatic mutations that could explain the observed clinical diversity. In EwS, chr8 gains are the second most common genetic alteration rendering EwS an ideal model to investigate the relevance of chr8 gains in an otherwise silent genomic context.

Here, we report that chr8 gain-driven gene expression patterns correlate with poor overall survival of EwS patients. This effect is predominantly mediated by increased expression of the translation initiation factor binding protein 4E-BP1 encoded by *EIF4EBP1* on chr8. High *EIF4EBP1* expression showed the strongest association with poor patient survival among all chr8-encoded genes and correlated with chr8 gains in EwS tumors. Similar findings were made in numerous entities of The Cancer Genome Atlas (TCGA). Integrated multi-omics profiling uncovered that 4E-BP1 orchestrates a pro-proliferative proteomic network. Consistently, silencing of 4E-BP1 in the EwS model reduced cell proliferation, clonogenicity, spheroidal growth *in vitro*, and tumorigenesis *in vivo*. Drug screens and functional assays revealed that high 4E-BP1 expression sensitizes for pharmacological CDK4/6 inhibition in preclinical models.

Collectively, we establish chr8 gains and high 4E-BP1 expression as prognostic biomarkers in EwS and demonstrate that their association with patient outcome is primarily mediated by 4E-BP1 orchestrating a pro-proliferative proteomic network sensitizing EwS for CDK4/6 inhibitors. Our data suggest that testing for chr8 gains may improve risk-stratification and therapeutic management in EwS and other cancers.

## INTRODUCTION

Aneuploidy is common in cancer cells and plays an important functional role in their pathophysiology^1–3^. Copy number alterations of chr8, especially chr8 gains, are observed in numerous cancer entities, including EwS, acute/chronic myeloid leukemia, gastric cancer, myxoid liposarcoma, pediatric undifferentiated sarcoma, clear cell sarcoma, and malignant peripheral nerve sheath tumors^2,4–11^. However, the functional and clinical role of chr8 gains remains to be clarified. In the context of precision oncology, understanding the role of specific chromosomal gains and losses as one major source of inter-tumor heterogeneity is important for the development of novel personalized diagnostic and therapeutic approaches.

EwS is a malignant bone- and soft tissue-related tumor, primarily occurring in children, adolescents, and young adults^12^. It is characterized by a low number of recurrent somatic mutations and is driven by chromosomal translocations generating pathognomonic *FET::ETS* fusions, with *EWSR1::FLI1* being the most common (present in 85% of cases), encoding aberrant chimeric transcription factors^12^. Genetic variants in polymorphic enhancer-like DNA binding sites of EWSR1::FLI1 were shown to account for inter-individual heterogeneity in EwS susceptibility, tumor growth, clinical course, and treatment response^13–15^. Secondary somatic mutations in *STAG2* and *TP53* occur in approximately 20 and 5% of EwS patients^16–18^, respectively. However, little is known about other even more common recurrent alterations, such as chromosomal gains and/or losses, and their impact on inter-individual tumor heterogeneity.

Chr8 gain is present in approximately 50% of EwS cases, often in form of chr8 trisomy, making it the second most frequently observed recurrent somatic alteration in EwS following *FET::ETS* fusions^16–22^. Previous studies focused solely on specific correlations regarding the role of (partial) chr8 gains in EwS^16–19,21,23–28^, and suggested that chr8 gains may be an early event in EwS tumorigenesis^29^. However, the precise functional and clinical impact of whole chr8 gains in EwS remains unclear. EwS serves as an ideal model to investigate the role of chr8 gain in cancer, given that EwS exhibit a ‘silent’ genome, where chr8 gains occur in an oligomutated genomic context^12^.

Therefore, the present study aimed to investigate the possible association between whole chr8 gains and tumor progression in the EwS model and to identify the most clinically relevant genes located on chr8 that may functionally contribute to inter-individual variability in patient outcomes. Following an integrative functional genomics approach, we have identified the eukaryotic translation initiation factor 4E binding protein 1 (EIF4EBP1, alias 4E-BP1) as the most promising chr8 candidate gene. It is outstandingly associated with unfavorable patient outcome compared to all other captured genes located on chr8 and even across the entire EwS transcriptome. 4E-BP1 functions downstream of its inactivating kinase complex, mTORC1 (mammalian target of rapamycin complex 1), and is a key effector of the mTORC1 signaling pathway^30,31^. 4E-BP1 belongs to a family of eIF4E-binding proteins that enable mTORC1 to adjust mRNA translation rates in response to various stimuli by modulating the assembly of the 48S translation pre-initiation complex. However, its role in tumor initiation/progression has not yet been defined^30^. We demonstrate that overexpression of *EIF4EBP1* is mediated by chr8 gain in primary EwS tumors. Furthermore, its RNAi-mediated knockdown in cell line models reduces EwS growth *in vitro* and *in vivo,* by influencing a pro-proliferative proteomic network. Thus, we establish an association between chr8 gain and tumor progression, mediated by 4E-BP1 in EwS. Drug screens and drug sensitivity assays *in vitro* and *in vivo* revealed that high 4E-BP1 expression sensitizes cells to targeted CDK4/6 inhibitor treatment with the FDA-approved drugs Palbociclib and Ribociclib. This discovery offers a new therapeutic strategy for tumors with chr8 amplification and 4E-BP1 overexpression.

## RESULTS

### Chromosome 8 gain drives overexpression of the clinically relevant translation initiation factor 4E-BP1 in EwS

To gain initial insights into whether chr8 gain mediates poor patient outcomes in EwS, we analyzed a cohort of 196 EwS samples for which matched microarray gene expression data and clinical data were available (henceforth referred to as ‘Cohort 1’). We used a chr8 gene expression signature as a surrogate model for factual genomic chr8 gain and performed a single-sample Gene Set Enrichment Analysis (ssGSEA) followed by hierarchical clustering, assigning each patient to either a high or low chr8 gene expression signature group (**Figure 1a**). To validate our approach, we first identified the differentially expressed genes (DEGs) between the inferred chr8 high and low clusters. We then performed a Position Related Data Analysis (PREDA) to map the respective DEGs to chromosomal positions, demonstrating that the vast majority of DEGs map to chr8 (**Supplementary Figure 1a**). Secondly, we applied our approach to RNA-sequencing data from an independent cohort of 100 EwS tumors (henceforth referred to as ‘Cohort 2’) and compared the chr8 signature enrichment clustering with matched factual chr8 genome-wide copy-number variation (CNV) status (inferred from DNA methylation arrays). This analysis showed that clustering based on the chr8 gene expression signature enrichment accurately indicates the presence of chr8 gain (**Supplementary Figure 1b**).

**Figure 1:**
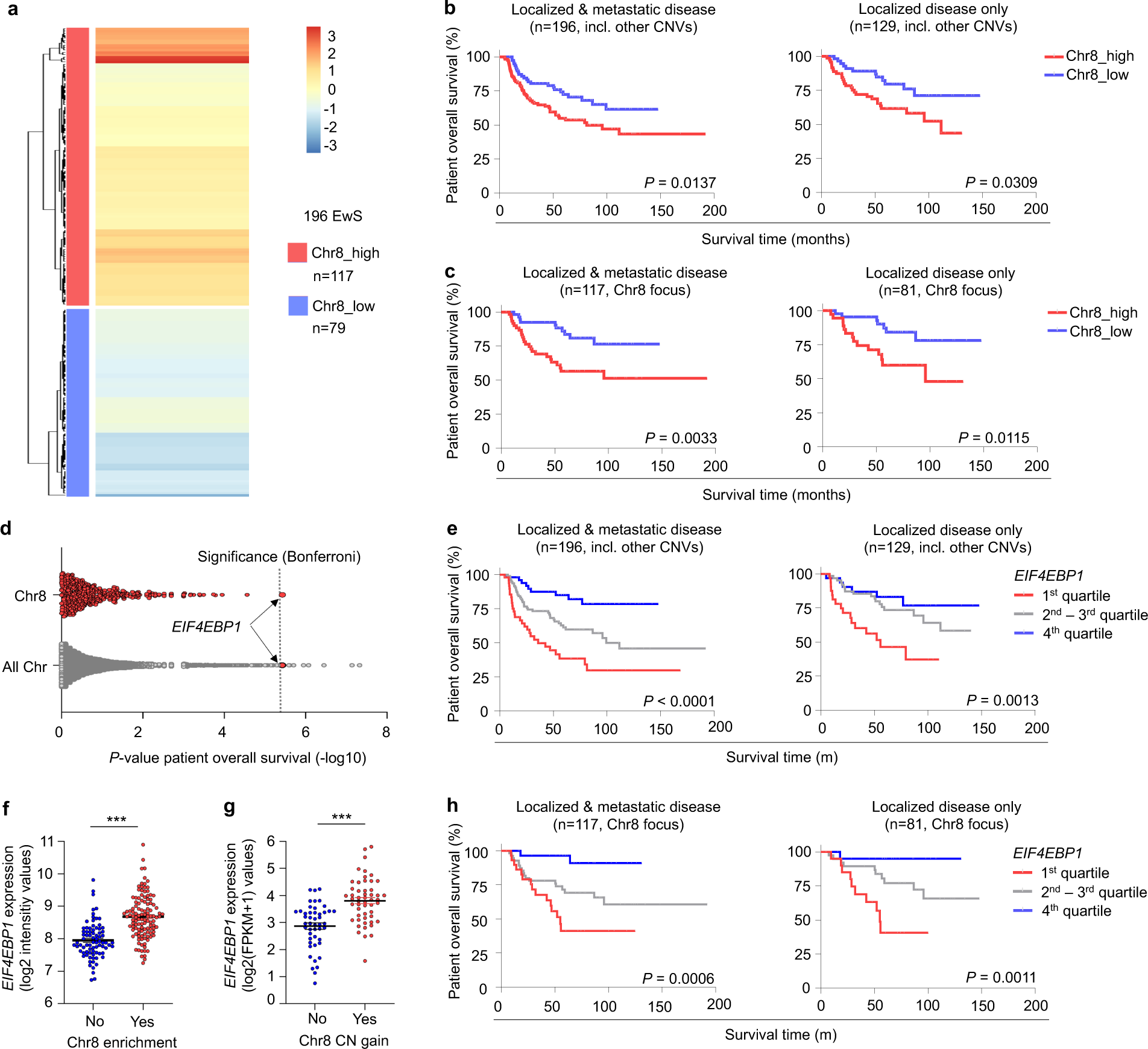
Chromosome 8 gain drives overexpression of the clinically relevant translation initiation factor 4E-BP1 in EwS. **a** Heat map showing 196 EwS patient tumors (Cohort 1) clustered according to an inferred chr8 gene expression signature, as assessed by single-sample Gene Set Enrichment Analysis (ssGSEA) followed by hierarchical clustering, assigning each sample into either a high or low chr8 signature gene expression group. **b** Kaplan-Meier overall survival analysis of 196 EwS patients (Cohort 1) stratified into either a high or low chr8 signature enrichment group as described in (a). Kaplan-Meier plots are shown separately either for patients with localized & metastatic disease (n=196, left) or exclusively for patients with localized disease (n=129, right). *P-*values determined by Mantel-Haenszel test. **c** Kaplan-Meier overall survival analysis of 117 EwS patients (Cohort 1, Chr8 focus) stratified into either a high or low chr8 signature enrichment group as described in (a) but excluding samples with other inferred recurrent CNVs. Kaplan-Meier plots are shown separately either for patients with localized & metastatic disease (n=117, left) or exclusively for patients with localized disease (n=81, right). *P-*values determined by Mantel-Haenszel test. **d** Overall survival batch analysis as assessed for every gene covered in transcriptomic profiling of 196 EwS patients (Cohort 1) using Mantel-Haenszel statistics. Chr8-located genes were additionally depicted separately. The dashed line indicates the Bonferroni-adjusted *P-*value threshold for significance. **e** Kaplan-Meier overall survival analysis of 196 EwS patients (Cohort 1) stratified by quartile *EIF4EBP1* expression. Kaplan-Meier plots are shown separately either for patients with localized & metastatic disease (n=196, left) or exclusively patients with localized disease (n=129, right). *P-*values determined by Mantel-Haenszel test. **f** *EIF4EBP1* expression as measured by microarray profiling in 196 EwS patient tumors (Cohort 1) stratified into either a high or low chr8 signature enrichment group as described in (a). *P-*values determined by two-tailed Mann-Whitney test, horizontal bars represent means and whiskers represent the SEM. \*\*\**P* < 0.001. **g** *EIF4EBP1* expression as measured by RNA-seq in 100 EwS patient tumors (Cohort 2) depending on the presence of chr8 gain as determined by methylation array. *P-*values determined by two-tailed Mann-Whitney test, horizontal bars represent means and whiskers represent the SEM. \*\*\**P* < 0.001. **h** Kaplan-Meier overall survival analysis of 117 EwS patients (Cohort 1, Chr8 focus as in (c)) stratified by quartile *EIF4EBP1* expression. Kaplan-Meier plots are shown separately either for patients with localized & metastatic disease (n=117, left) or exclusively patients with localized disease (n=81, right). *P-*values determined by Mantel-Haenszel test.

Kaplan-Meier analysis in Cohort 1 revealed that a high chr8 gene expression signature was associated with shorter overall EwS patient survival (*P*=0.0137, **Figure 1b**). Strikingly, this association remained significant (*P*=0.0309, **Figure 1b**) even when only considering patients with localized disease (i.e., without evidence for metastasis at diagnosis), indicating that chr8 gain is functionally involved in mediating an unfavorable disease phenotype. In support of this hypothesis, it is intriguing that while chr8 gain is only found in approximately 50% of primary tumors, around 80% of EwS cell lines, which are expected to be derived from highly aggressive tumor clones, exhibit chr8 gains (mostly trisomies)^16–19,21,24–28^. Since previous studies have reported that chr8 gains can co-occur with other recurrent chromosomal gains and losses that may have an effect on patient overall survival^16–18,27^, such as chr1q gains, chr12 gains, and 16q loss, we reassessed our Cohort 1 now only focusing on those patients that show a predicted exclusive chr8 gain or none of the above-mentioned CNVs. As shown in **Figure 1c**, focusing on exclusively chr8 gained samples yielded even a better patient-stratification regarding overall survival in both localized disease and the entire sub-cohort (*P*=0.0115 and *P*=0.0033 respectively). Together, these findings suggest that genes located on chr8 contribute to aggressive cellular behavior and disease progression in EwS. Previous reports suggested that *MYC* located on chr8 may mediate the effect of chr8 gains on patient outcome in EwS and other undifferentiated sarcomas^10,32^. However, in our large EwS Cohort 1, *MYC* expression was not significantly associated with overall patient survival (*P*=0.689, **Supplementary Figure 1c**, **Supplementary Table 1**). Similarly, the chr8 gene *RAD21*, previously reported to promote EwS tumorigenicity by mitigating EWSR1::FLI1-induced replication stress^23^, did not show a significant association with overall EwS patient survival (*P*=0.174; **Supplementary Figure 1c, Supplementary Table 1**). These findings suggest that the mechanisms underlying the association of chr8 gain with EwS aggressiveness are more complex than previously anticipated.

Thus, to identify genes on chr8 strongly associated with overall survival of EwS patients, we performed a batch-analysis within Cohort 1 calculating *P*-values for the association with overall patient survival for every gene represented on the microarray using our custom code software GenEx applying Mantel-Haenszel statistics (**Supplementary Table 1**). Analysis of all covered genes located on chr8 revealed that the expression of *EIF4EBP1* showed the strongest association with overall patient survival and, more specifically, that high expression of *EIF4EBP1* significantly correlated with unfavorable overall survival of EwS patients (nominal *P*<0.0001, Bonferroni-adjusted *P*=0.049, **Figure 1d,e**, **Supplementary Table 2**). High *EIF4EBP1* expression remained significantly associated with poor overall survival even when considering only patients with localized disease (*P*=0.0013, **Figure 1e**). Additionally, *EIF4EBP1* ranked within the top 15 survival associated genes genome-wide (**Figure 1d, Supplementary Table 1**). These results are in consistence with the association of chr8 gain with poor overall survival in EwS patients (**Figure 1b,c**) as well as with previous research that has linked chr8p, where *EIF4EBP1* is located, with EwS relapse^32,33^. Furthermore, *EIF4EBP1* expression is significantly correlated with high ssGSEA enrichment scores for chr8 gene expression in Cohort 1 (*P*<0.001, Pearsońs r = 0.47, Coheńs d = 1.19). This suggests that a significant part of the negative prognostic effect of the high chr8 gene expression signature can be attributed to high *EIF4EBP1* expression. Accordingly, the predicted chr8 gain is significantly associated with elevated *EIF4EBP1* expression levels in this cohort (P<0.001; **Figure 1f**), which was fully confirmed in the independent Cohort 2 with chr8 status detected at the DNA level (*P*<0.001; **Figure 1g**). Similar to our analyses shown in **Figure 1c**, we reanalyzed our survival data from Cohort 1 now only focusing on solely chr8 gained samples versus samples without any recurrent chromosomal gain/loss, which fully confirmed the prognostic role of *EIF4EBP1* in EwS patients (**Figure 1h**). Interestingly, DEG analysis of Cohort 1 comparing chr8 high and low gene expression revealed that *EIF4EBP1* is distinctively differentially upregulated within genes of the mTOR signaling pathway (**Supplementary Figure 1d**), indicating that 4E-BP1 has a distinct clinical and functional role within the mTOR signaling pathway in EwS.

To evaluate the potential clinical and functional significance of chr8 gain and *EIF4EBP1* expression in other cancer entities besides EwS, we analyzed CNV data from DNA methylation arrays of The Cancer Genome Atlas (TCGA). Our analysis revealed that numerous cancer entities exhibit chr8 gains (specifically, 8 out of 32 identified entities exhibit chr8 gains in more than 10% of cases) (**Supplementary Table 3**). Additionally, chr8 gain and high *EIF4EBP1* expression are associated with unfavorable patient survival in several other cancer entities (chr8 gain in 4 and high *EIF4EBP1* expression in 14 out of 32 identified entities) (**Supplementary Table 3**). These include hepatocellular carcinoma, renal papillary cell carcinoma, lower-grade glioma, and thymoma (**Supplementary Table 3**).

Collectively, these results indicate that chr8 gain contributes to unfavorable outcomes of EwS patients and identify 4E-BP1 as a potential driver of EwS aggressiveness encoded on chr8.

### 4E-BP1 drives a multifunctional proliferation-associated proteomic network

Contrary to our findings that high *EIF4EBP1* levels significantly correlated with worse patient outcome (**Figure 1d,e,h**), a recent report has suggested that 4E-BP1 may act as a tumor suppressor in EwS^34^. However, this conclusion was based on observations upon supraphysiological, ectopic overexpression of a phospho-mutant (and thus functionally inactive) 4E-BP1 protein in two EwS cell lines (EW8 and TC-71)^34^. The role of 4E-BP1 in cancer is complex and strongly depends on the cellular context and its precise phosphorylation status^30^. Therefore, to obtain a more comprehensive understanding of 4E-BP1 in EwS, we first integrated results of pre-ranked fast Gene Set Enrichment Analyses (fGSEA). We conducted fGSEAs based on Pearson’s correlation coefficients between the mRNA expression levels of *EIF4EBP1* and every other gene represented in the respective datasets of Cohort 1 and 2 (**Supplementary Table 4,5**). Additionally, we carried out a third fGSEA based on gene expression fold-changes (FCs) between tumors with and without detected chr8 gain in Cohort 2 (**Supplementary Table 6**). The overlap between all three fGSEAs consisted predominantly of proliferation-associated gene sets (**Figure 2a,b, Supplementary Table 4-6**). These transcriptomic data from EwS patients pointed toward a role of 4E-BP1 in the regulation of EwS cell proliferation and strongly supported the potential role of 4E-BP1 as a major mediator of chr8 gain-driven poor prognosis in EwS.

**Figure 2:**
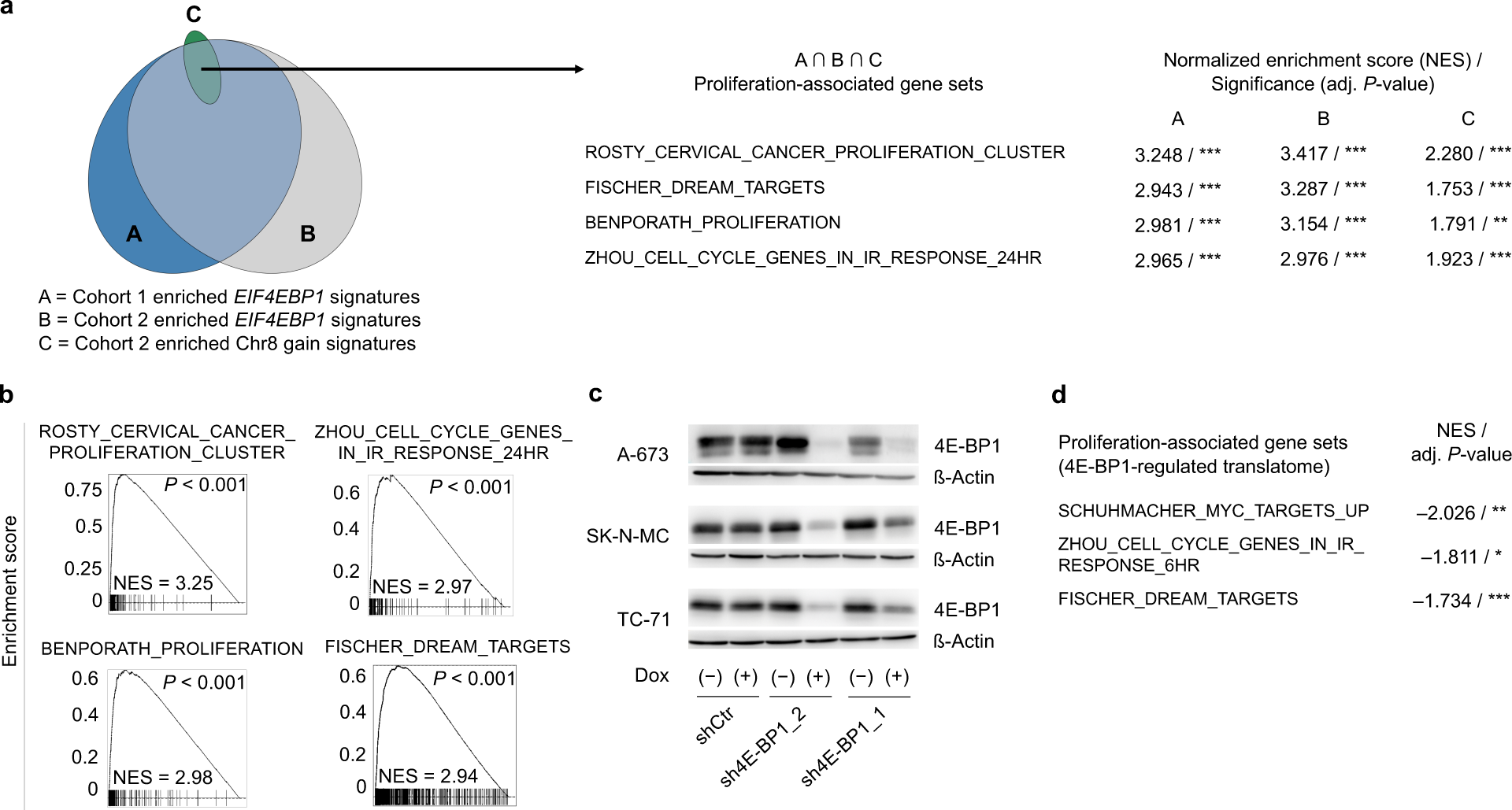
4E-BP1 drives a multifunctional proliferation-associated proteomic network. **a** Area-proportional Venn diagram of gene sets enriched with *EIF4EBP1* expression in Cohort 1 (A) and 2 (B) as well as with chr8 gain in Cohort 2 (C) as determined by fGSEA. Exemplary gene sets representing a proliferation-associated enrichment signature in the overlap between A, B, and C are shown with respective normalized enrichment scores (NES) and significance levels. \*\*\**P* < 0.001 **b** fGSEA enrichment plots of exemplary gene sets displayed in (a). **c** Representative western blots in A-673, SK-N-MC, and TC-71 cells containing either Dox-inducible specific shRNA constructs directed against *EIF4EBP1* (sh4E-BP1_1 or sh4E-BP1_2) or a non-targeting shControl (shCtr). Cells were grown either with or without Dox for 96 h. ß-actin served as a loading control. **d** Gene sets negatively enriched upon 4E-BP1 knockdown on protein level, as determined by fGSEA using integrated mass spectrometry and microarray protein/gene expression data as an input. Exemplary gene sets representing a proliferation-associated enrichment signature are shown with respective normalized enrichment scores (NES) and significance levels (\*\*\**P* < 0.001, \*\**P* < 0.01, \**P* < 0.05).

To further explore this hypothesis, we generated an *in vitro* 4E-BP1 knockdown model in three EwS cell lines with relatively high *EIF4EBP1* baseline expression levels (A-673, SK-N-MC, and TC-71) (**Supplementary Figure 2a**). Notably, two of the selected cell lines (SK-N-MC and TC-71) exhibit a chr8 gain^26^. To that end, we transduced EwS cell lines with a lentivirus containing a vector-system (pLKO Tet-on) with doxycycline (Dox)-inducible shRNAs, specifically directed against *EIF4EBP1* (sh4E-BP1_1 or sh4E-BP1_2) or a non-targeting control shRNA (shCtr). Both targeted shRNAs effectively silenced *EIF4EBP1* mRNA expression, resulting in a strong knockdown of *EIF4EBP1* mRNA levels (**Supplementary Figure 2b**) and protein levels (**Figure 2c, Supplementary Figure 2c**). This is consistent with a strong correlation between (EIF)4E-BP1 mRNA and protein levels in human cells, as evidenced by the analysis of Cancer Dependency Map (DepMap) gene expression and corresponding protein array data (*n*=887 cancer cell lines, *r_Pearson_*=0.68, *P*=5.2×10^-22^). Western blot experiments demonstrated that knockdown of 4E-BP1 led to a consistent loss of its phosphorylated form (Ser65) in all EwS cell lines in similar manner as total 4E-BP1 (**Supplementary Figure 2d**). Therefore, our EwS 4E-BP1 knockdown models are well suited to study the functional consequences of its inactivation.

As 4E-BP1 regulates mRNA translation initiation by binding to the translation initiation factor eIF4E and thereby modifying overall and selective translation rates^30^, we asked whether functional interference with 4E-BP1 might also affect proliferation-related translational signatures.

To identify proteins regulated by 4E-BP1 exclusively at the translational level, we combined mass spectrometry (MS) based proteomic profiling of newly synthesized proteins with parallel transcriptome profiling by gene expression microarrays. To this end, we silenced 4E-BP1 in three EwS cell lines (A-673, SK-N-MC, and TC-71) and pulsed them with stable isotope labeled amino acids in cell culture (SILAC) medium and the methionine analog L-Azidohomoalanine (AHA) for 6 h. We identified 9,508 proteins through MS analysis, of which 4,335 common proteins across all cell lines and constructs with at least one value per replicate group were used for downstream analyses. Our parallel microarray analyses captured 12,056 stably expressed genes across all three cell lines. Following further filtering steps (see methods section), we identified 1332 differentially expressed proteins upon 4E-BP1 knockdown (adj. *P* value < 0.05), which were not regulated by 4E-BP1 at the mRNA level across all three cell lines (**Supplementary Table 7**). Preranked fGSEA analysis on proteins not regulated at the mRNA level, and therewith most likely directly differentially regulated by 4E-BP1, identified again a strong enrichment of proliferation-associated gene sets (**Figure 2d, Supplementary Table 8**) consistent with fGSEA results from patient gene expression data as shown in **Figure 2a,b**.

Our *in silico* analyses of patient data at the mRNA level and functional *in vitro* analyses at the protein level collectively indicate that 4E-BP1 is linked to accelerated proliferation of EwS cells, suggesting a potential role as an oncogene in EwS.

### 4E-BP1 promotes proliferation and tumorigenicity of EwS cells

To confirm the pro-proliferative and oncogenic role of 4E-BP1 in EwS, we conducted various functional *in vitro* and *in vivo* assays. We found that knockdown of 4E-BP1 for 96 h significantly inhibited cell proliferation in all three cell lines (**Figure 3a**). The anti-proliferative effect of 4E-BP1 knockdown appeared to be independent of cell death as Trypan-Blue-exclusion assays did not consistently show a significant effect of 4E-BP1 knockdown on cell death across all cell lines and shRNAs (**Supplementary Figure 3a**). Prolonged 4E-BP1 knockdown (10–14 d) significantly reduced both 2D clonogenic and 3D anchorage-independent growth of EwS cells (**Figure 3b,c**). Such effects were not observed in shCtr cells (**Figure 3b,c**). Similarly, knockdown of 4E-BP1 in subcutaneously xenotransplanted cells significantly reduced tumor growth *in vivo* (**Figure 3d**, **Supplementary Figure 3b**). Consistent with our *in vitro* results, this phenotype was linked to a significantly diminished mitotic cell count, as revealed by histologic assessment of the respective xenografts (**Figure 3e**, **Supplementary Figure 3c**). No difference in tumor necrosis was observed between xenografts with or without 4E-BP1 knockdown (**Supplementary Figure 3d**). To validate the effect of 4E-BP1 in an orthotopic xenograft model, we xenografted TC-71 cells transduced with an inducible *EIF4EBP1* targeting shRNA construct (sh4E-BP1_2) into the proximal tibia of NSG mice, which were subsequently treated with or without Dox. Similar to our subcutaneous xenograft model, the tumor burden in orthotopic EwS xenografts decreased upon Dox-induced knockdown of 4E-BP1 (**Figure 3f**).

**Figure 3:**
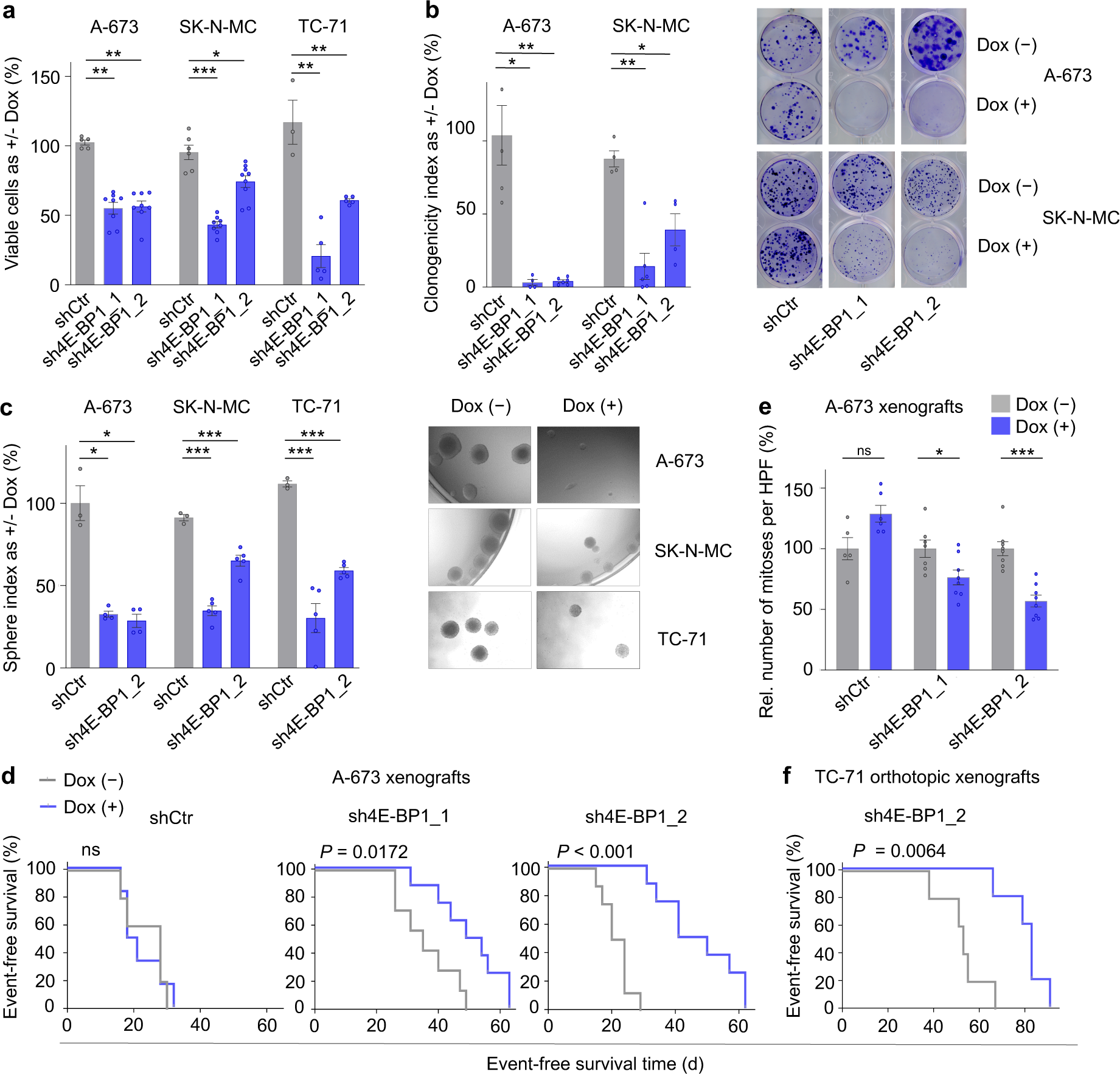
RNAi-mediated knockdown of 4E-BP1 inhibits EwS growth. **a** Relative viable cell count of A-673, SK-N-MC, and TC-71 cells containing either Dox-inducible specific shRNA constructs directed against *EIF4EBP1* (sh4E-BP1_1 or sh4E-BP1_2) or a non-targeting shControl (shCtr). Cells were grown either with or without Dox for 120 h. Horizontal bars represent means, and whiskers represent the SEM, n≥4 biologically independent experiments. **b** Relative colony number of A-673 and SK-N-MC cells containing either Dox-inducible specific shRNA constructs directed against *EIF4EBP1* (sh4E-BP1_1 or sh4E-BP1_2) or a non-targeting shControl (shCtr). Cells were grown either with or without Dox for 8–14 d. Horizontal bars represent means, and whiskers the SEM, n≥4 biologically independent experiments. Representative images of colony formation are shown on the right. **c** Sphere formation in A-673, SK-N-MC, and TC-71 cells containing shRNA constructs directed against *EIF4EBP1* (sh4E-BP1_1 or sh4E-BP1_2) or a non-targeting shControl (shCtr) treated with or without Dox for 8–14 d. Horizontal bars represent means, and whiskers represent the SEM, n≥3 biologically independent experiments. *P*-values determined by two-tailed unpaired t-test with Welch’s correction. Representative images of spheres are shown on the right. **d** Kaplan-Meier analysis of event-free survival of NSG mice xenografted with A-673 cells containing either Dox-inducible specific shRNA constructs directed against *EIF4EBP1* (sh4E-BP1_1 or sh4E-BP1_2) or a non-targeting shControl (shCtr). Once tumors were palpable, mice were randomized and treated with either vehicle (–) or Dox (+), n≥5 animals per condition. An ‘event’ was recorded when tumors reached a size maximum of 15 mm in one dimension. *P-*values determined via Mantel-Haenszel test. **e** Quantification of mitoses in HE-stained slides of xenografts described in (d). Five high-power fields (HPF) were counted per sample. Horizontal bars represent means, and whiskers represent the SEM, n≥4 samples per condition. **f** Kaplan-Meier analysis of event-free survival of NSG mice orthotopically xenografted into the proximal tibia with TC-71 cells containing a Dox-inducible specific shRNA construct directed against *EIF4EBP1* (sh4E-BP1_2). One day after injection of the cells, mice were randomized and treated with either vehicle (–) or Dox (+), n = 5 animals per condition. An ‘event’ is recorded when the mice exhibited signs of limping at the injected leg. *P-*values determined via Mantel-Haenszel test. \*\*\**P* < 0.001, \*\**P* < 0.01, \**P* < 0.05, ns = not significant; *P*-values determined via two-tailed Mann-Whitney test if not otherwise specified.

In summary and in conjunction with our integrative clinical and *in silico* analyses from patient tumors and cell line models (**Figure 1 and 2)**, these results generated *in vitro* and *in vivo* provide strong evidence that 4E-BP1 acts as an oncogene in EwS.

### High 4E-BP1 expression sensitizes for CDK4/6 inhibitor treatment

To identify therapeutic vulnerabilities in EwS with high 4E-BP1 expression, we conducted drug screens on 3D-spheroids of A-673 EwS cells with/without knockdown of 4E-BP1. Ribociclib, an FDA-approved CDK4/6 inhibitor^35–38^, was the top hit demonstrating differential sensitivity in 4E-BP1 high expressing cells (**Supplementary Figure 4a**). The presented data align with the published gene-dependency data of DepMap project, indicating a significant and selective dependency of EwS cell lines on CDK4 expression compared to non-EwS cell lines (**Supplementary Figure 4b**). We validated these findings in 2D culture experiments using A-673 EwS cells with/without knockdown of 4E-BP1, additionally using a second FDA-approved CDK4/6 inhibitor, Palbociclib (**Figure 4a**)^35–38^. Interestingly, EwS cell lines with high endogenous 4E-BP1 expression (A-673 and TC-71) showed greater sensitivity to Palbociclib and Ribociclib than cell lines with low endogenous 4E-BP1 expression (EW-22 and CHLA-10) (**Figure 4b**).

**Figure 4:**
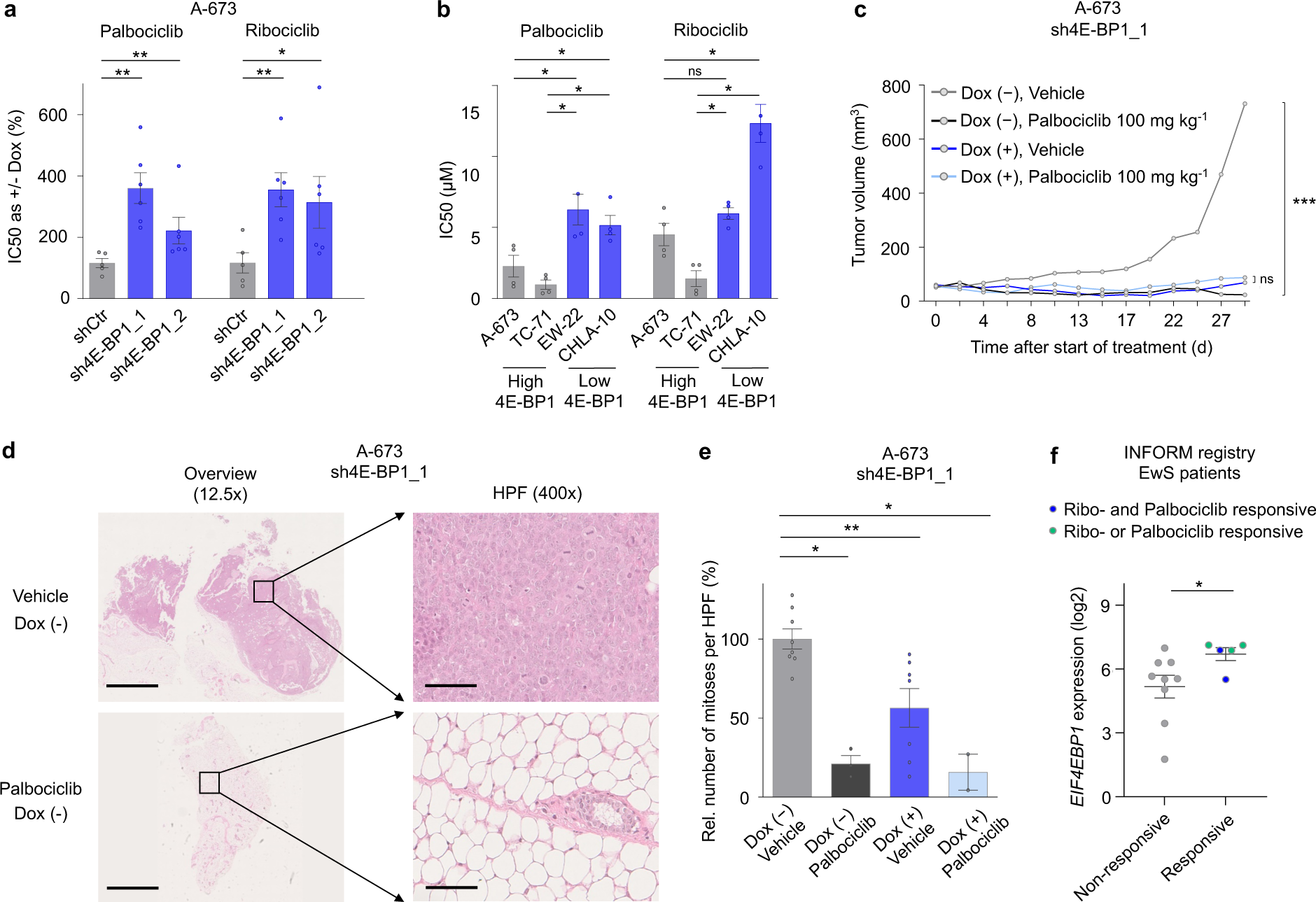
High 4E-BP1 expression sensitizes to targeted CDK4/6 inhibitor treatment with Palbociclib and Ribociclib. **a** IC50 analysis of CDK4/6 inhibitors Palbociclib and Ribociclib in A-673 cells containing either DOX-inducible specific shRNAs directed against 4E-BP1 (sh4E-BP1_1, sh4E-BP1_2) or a non-targeting a non-targeting shControl (shCtr) as measured by resazurin colorimetry. Cells were treated with/without Dox as well as with serial dilutions of respective inhibitors. Horizontal bars represent means, and whiskers represent the SEM, n ≥ 5 biologically independent experiments. **b** IC50 analysis of CDK4/6 inhibitors Palbociclib and Ribociclib in EwS cells with high (A-673, TC-71) and low (EW-22, CHLA-10) endogenous 4E-BP1 expression as measured by resazurin colorimetry. Cells were treated with serial dilutions of respective inhibitors. Horizontal bars represent means, and whiskers represent the SEM, n≥3 biologically independent experiments. **c** NSG mice xenografted with A-673 EwS cells containing a Dox-inducible sh4E-BP1 construct, treated with/without Dox and either vehicle or Palbociclib in a dose of 100 mg/kg. Mice were randomized to the treatment groups when tumors were palpable. For each condition the mean tumor volume and SEM of 4–6 mice over the time of treatment are shown. **d** Representative HE stained micrographs of A673/sh4E-BP1 xenografts (Dox (–)) treated with either vehicle or Palbociclib as described in (c) (shown as an overview with 12.5*×* magnification and as a high-power field (HPF) in 400*×* magnification). Scale bar is 2.5 mm (12.5*×*) and 100 µm (400*×*). **e** Quantification of mitoses in micrographs of xenografts described in (c). Horizontal bars represent means, and whiskers represent the SEM, n≥2 samples per condition. **f** *EIF4EBP1* gene expression data from 14 EwS patient tumors treated within the INFORM registry stratified according to matched Palbociclib/Ribociclib drug sensitivity data from 3D tumor cell cultures into a CDK4/6 inhibitor non-responsive and responsive group. \*\*\**P* < 0.001, \*\**P* < 0.01, \**P* < 0.05, ns = not significant; *P*-values determined via two-tailed Mann-Whitney test if not otherwise specified.

Next, we conducted xenograft experiments by transplanting A673 EwS cells subcutaneously into the flanks of NSG mice, treated with/without Dox and with/without Palbociclib. Xenografts with 4E-BP1 knockdown and xenografts with Palbociclib treatment similarly exhibited a very strong reduction of tumor growth, which correlated with a strong decrease in histologically assessable viable tumor burden as compared to respective xenografts without 4E-BP1 knockdown or Palbociclib treatment (**Figure 4c,d, Supplementary Table 9**). Consistently, xenografts of mice with 4E-BP1 knockdown or treatment with Palbociclib, for which histological material was obtainable, showed a lower number of mitoses per high-power field (**Figure 4e**). However, since the very strong growth-inhibitory effects of either 4E-BP1 knockdown or Palbociclib treatment alone precluded the assessment of a potential differential effect of 4E-BP1 expression on sensitivity toward Palbociclib in this model (**Figure 4c,d, Supplementary Table 9**), we turned for further validation to patient-derived real-world data. To this end, we analyzed gene expression and 3D drug sensitivity data from 14 EwS patient-derived short-term cultures treated with Palbociclib or Ribociclib in the context of the Individualized Therapy For Relapsed Malignancies in Childhood (INFORM) registry^39^. Strikingly, we found that high *EIF4EBP1* expression is indeed associated with higher sensitivity toward CDK4/6 inhibitor treatment (**Figure 4f**).

In summary, these results suggest that 4E-BP1 may serve as a valuable predictive biomarker for clinical effectiveness of CDK4/6 inhibitor treatment.

## DISCUSSION

The data presented herein in the EwS model establish chr8 gain as an unfavorable prognostic factor, primarily mediated through the overexpression of the translation initiation factor 4E-BP1, which guides pro-proliferative proteomic signatures and sensitizes cells to targeted CDK4/6 inhibitor treatment.

In precision oncology, it is crucial to decipher mechanisms underlying inter-tumoral heterogeneity to refine diagnostic and therapeutic algorithms^14,40,41^. In this context, the identification of chr8 gain as a prognostic factor emphasizes the relevance of cytogenetic testing, which may help stratify patients into prognostic and/or therapeutic subgroups. Although chr8 trisomies are observed in approximately 50% of EwS patients, so far, only trends or moderate associations between whole chr8 gain (i.e., trisomies) and poor patient outcome have been observed^21,23–25,32,42,43^. Our data provide evidence of a significant association between a high chr8 gene expression signature and poor overall survival of EwS patients (**Figure 1**). Importantly, this association remains statistically significant even when only considering patients with localized disease (**Figure 1**). Therefore, chr8 gain, as assessed by cytogenetic testing or fluorescence in situ hybridization (FISH) or a high chr8 gene expression signature score (i.e., as assessed by ssGSEA), might serve as a prognostic biomarker for poor overall patient survival. Consequently, it could be particularly useful for stratifying patients with localized disease into different treatment groups.

Our results are consistent with data from other cancer entities that have shown a prognostic/predictive value of chr8 gain, as for example in acute myeloid leukemia^6^ and chronic myeloid leukemia^4,5^. Interestingly, chr8 gain is also observed in several specific other sarcoma entities, including myxoid liposarcoma^9^, clear cell sarcoma^8^, and pediatric undifferentiated sarcomas^10^. Partial gains or losses of chromosome 8 have been described in a broad range of cancer entities, such as prostate, lung, hepatocellular, and renal cell carcinoma^44–48^. In contrast to the reported data from EwS, in other cancer entities, chr8p losses are most frequently described to be associated with unfavorable clinical parameters^46,48,49^. In the case of chr8q, gains are most frequently described as having tumor promoting functions due to resulting *MYC* amplification^44,47,48,50^. However, in our patient cohort, a clinical association between *MYC* expression and overall survival was not evident (**Supplementary Figure 1c**, **Supplementary Table 1**). Furthermore, other possible mechanisms of *EIF4EBP1* upregulation, apart from chr8 gains, have been described across cancer entities. For example, direct upregulation driven by differential transcription factors may account for high 4E-BP1 expression levels in individual tumors^14,51^.

We demonstrate that high 4E-BP1 expression levels sensitize EwS cells to CDK4/6 inhibitor treatment with Palbociclib and Ribociclib (**Figure 4**). These drugs are FDA-approved for the treatment of hormone receptor positive, human epidermal growth factor receptor 2 (EGFR2, alias HER2) negative advanced or metastatic breast cancer, used in combination with an aromatase inhibitor in postmenopausal women^35–38^. In addition to their prognostic value in EwS (**Figure 1**), chr8 gain and 4E-BP1 expression might serve as predictive markers to subject EwS patients into a CDK4/6-inibitor sensitive and non-sensitive groups. Such tailored stratification of patients into specific targeted treatment groups with already FDA-approved drugs could significantly and promptly improve EwS patient outcome within the context of precision oncology. Notably, preclinical studies have shown that IGF-1R activation can mediated CDK4/6 inhibitor resistance^52^. As a result, a phase II clinical trial is currently investigating Palbociclib in combination with the IGF-1R inhibitor Ganitumab for patients with relapsed or refractory EwS^53^. However, this study did not include patient stratification based on predictive biomarkers. This gap might be addressed in future studies by incorporating predictive testing for chr8 gain and/or 4E-BP1 expression.

Collectively, our data suggest that chr8 gain plays an important prognostic role in EwS and that its functional effects on tumor progression are primarily driven by increased 4E-BP1 expression mediating a pro-proliferative phenotype. Since chr8 gain occurs in approximately 50% of EwS cases, our results indicate that this chromosomal aberration is a major source of inter-tumoral heterogeneity in EwS. Consequently, further cytogenetic testing of EwS might offer a refinement of clinical management within the context of precision oncology.

## MATERIALS AND METHODS

### Provenience of cell lines and cell culture conditions

Human EwS cell line A-673 (RRID: CVCL_0080) and human HEK293T cells (RRID: CVCL_0063) were purchased from American Type Culture Collection (ATCC). Human EwS cell line SK-N-MC (RRID: CVCL_0530) was provided by the German collection of Microorganisms and Cell cultures (DSMZ). Human EwS cell line TC-71 (RRID: CVCL_2213) was provided by the Children’s Oncology Group (COG). All cell lines were grown at 37 °C and 5% CO_2_ in humidified atmosphere in RPMI 1640 medium supplemented with stable glutamine (Biochrom), 10% tetracycline-free fetal calf serum (FCS) (Sigma-Aldrich), 100 U/ml penicillin (Biochrom), and 100 µg/ml streptomycin (Biochrom) was used to grow the cells. Cells were routinely examined by nested PCR for mycoplasma infection. Cell line purity and authenticity was confirmed by STR-profiling.

### Nucleotide extraction, reverse transcription, and quantitative real-time PCR (qRT-PCR)

Genomic DNA from human cell lines was extracted with the NucleoSpin Tissue kit (Macherey-Nagel), plasmid DNA was extracted from bacteria with the PureYield kit (Promega). RNA was extracted with the NucleoSpin II kit (Macherey-Nagel) and reverse-transcribed using the High-Capacity cDNA Reverse Transcription kit (Applied Biosystems). qRT-PCR reactions were performed using SYBR green Mastermix (Applied Biosystems) mixed with diluted cDNA (1:10) and 0.5 µM forward and reverse primer (total reaction volume 15 µl) on a BioRad Opus instrument and analyzed using BioRad CFX Manager 3.1 software. Gene expression values were calculated using the 2^-(ΔΔCt)^ method^54^ relative to the housekeeping gene *RPLP0* as an internal control. Oligonucleotides were purchased from MWG Eurofins Genomics and are listed in **Supplementary Table 10.** The thermal conditions for qRT-PCR were as follows: initialization (95 °C, 2 min) (1 cycle); denaturation (95 °C, 10 sec), annealing (60 °C, 10 sec), and extension (60 °C, 10 sec) (49 cycles); denaturation (95 °C, 30 sec), annealing (65 °C, 30 sec), extension (melting curve 65 °C increasing 0.5 °C every 5 sec until 95 °C) (1 cycle).

### Generation of doxycycline (Dox)-inducible shRNA constructs

Human EwS cell lines A-673, SK-N-MC, and TC-71 were transduced with lentiviral Tet-pLKO-puro all-in-one vector system (plasmid #21915, Addgene) containing a puromycin-resistance cassette, and a tet-responsive element for Dox-inducible expression of shRNAs against *EIF4EBP1* (sh4E-BP1_1 or sh4E-BP1_2) or a non-targeting control shRNA (shCtr). Sequences of the used shRNAs are listed in **Supplementary Table 10**. Dox-inducible vectors were generated according to a publicly available protocol^55^ using In-Fusion HD Cloning Kit (Clontech). Vectors were amplified in Stellar Competent Cells (Clontech) and respective integrated shRNA was verified by Sanger sequencing. The used sequencing primer is listed in **Supplementary Table 10**. Lentiviral particles were generated in HEK293T cells. Virus-containing supernatant was collected to infect the human EwS cell lines. Successfully transduced cells were selected with 1.5 µg/ml puromycin (InvivoGen). The shRNA expression for *EIF4EBP1* knockdown or expression of a negative control shRNA in EwS cells was achieved by adding 0.5 µg/ml Dox every 48 h to the medium. Generated cell lines were designated as A-673/TR/shCtr, A-673/TR/sh4E-BP1_1, A-673/TR/sh4E-BP1_2, SK-N-MC/TR/shCtr, SK-N-MC/TR/sh4E-BP1_1, SK-N-MC/TR/sh4E-BP1_2, TC-71/TR/shCtr, TC-71/TR/sh4E-BP1_1, and TC-71/TR/sh4E-BP1_2.

### Western blot

A-673, TC-71, and SK-N-MC cells were treated for 96h with Dox to induce the *EIF4EBP1* knockdown. Whole cellular protein was extracted with RIPA buffer (Serva electrophoresis) containing protease inhibitor cocktail and phosphatase inhibitor cocktail (Roche). Western blots were performed following routine protocols^56^ and specific band detection was achieved by the use of rabbit monoclonal anti-4E-BP1 (clone Y329, 1:2,000, ab32024, Abcam) and rabbit polyclonal anti-p-4E-BP1 antibody (1:1,000, Ser65, #9451, Cell Signaling) and mouse monoclonal anti-ß-actin (1:5,000, A-5441, Sigma-Aldrich). Anti-rabbit IgG horseradish peroxidase coupled antibody (1:2,000, sc-516102, Santa Cruz Biotechnology) and anti-mouse IgG horseradish peroxidase coupled antibody (1:2,000, sc-2357, Santa Cruz Biotechnology) were used as secondary antibodies. Quantification of western blots was performed using ImageStudio Light software.

### Proliferation assays

Depending on the cell line, 30–70 **×** 10^3^ cells containing either a Dox-inducible non-targeting control shRNA or *EIF4EBP1*-targeting specific shRNAs were seeded in duplicate wells of a 6-well plate in 2 ml of growth medium. The cells were treated either with or without Dox (0.5 µg/ml; Sigma-Aldrich) for 120 h (medium was changed after 72 h including fresh Dox). Afterwards, cells of each treatment condition were harvested (including supernatant), stained with Trypan blue (Sigma-Aldrich), and viable as well as dead cells were counted in a standardized hemocytometer (C-chip, NanoEnTek). The assays were performed according to routine protocols^57^.

### Clonogenic growth assays (Colony forming assays)

Depending on the cell line, 500–1,000 cells containing either a Dox-inducible non-targeting control shRNA or *EIF4EBP1*-targeting specific shRNAs were seeded in triplicate wells of a 12-well plate in 2 ml of growth medium. Cells were grown with or without Dox (0.5 µg/ml) for 8–14 d depending on the cell line and afterwards stained with crystal violet (Sigma-Aldrich). Colony number and area was determined on scanned plates using Fiji (Image J)^58,59^. Clonogenicity index was calculated by multiplying the counted colonies with the corresponding colony volume.

### Spheroidal growth assays

Depending on the cell line, 1,500–4,000 cells containing either a Dox-inducible non-targeting control shRNA or *EIF4EBP1*-targeting specific shRNAs were seeded in Costar Ultra-low attachment plates (Corning) for 8–14 d in 150 µl growth medium. To maintain the 4E-BP1 knockdown, 10 µl of fresh medium with or without Dox was added every 48 h to the cells. At day 8–14, the wells were photographed and spheres larger than 500 µm in diameter were counted. The area was measured using ImageJ. The sphere volumes were calculated as follows: V = 4/3 × π × r^3^. The sphere index was calculated by multiplying the counted colonies with the corresponding colony volume.

### Drug screening assays (3D)

Drug screening experiments were performed essentially as previously described in Peterziel et al.^60^ using 384-well round bottom ultralow attachment spheroid microplates (# 3830, Corning) to allow the formation of three-dimensional spheroids.

Briefly, we tested a drug library of 75 drugs, mostly approved or in clinical trials, covering standard chemotherapeutic drugs, epigenetic modifiers, metabolic modifiers, kinase inhibitors, apoptotic modulators, and others^60–62^. Drug plates were ordered from FIMM High Throughput Biomedicine Unit (Institute for Molecular Medicine Finland HiLIFE, University of Helsinki, Finland) as ready-to-use assay plates and stored in an oxygen- and moisture-free environment (San Francisco StoragePod, Roylan Developments Ltd, Fetcham Leatherhead, UK) at room temperature until use. One drug plate set consisted of three plates. The concentration range of each drug covered five orders of magnitude with each condition tested in duplicate. Wells containing 100 µM benzethonium chloride (BztCl), 250 nM staurosporine (STS) (serving as a death control), and 0.1% DMSO were included as maximum, intermediate, and minimum effect controls, respectively. An STS concentration range of 0.1 to 1,000 nM served as a technical control (two replicates per plate). Per well, 25 µl of single-cell suspension was dispensed on ready-to-use plates with an 8-channel electronic Picus pipette (10 to 300 µl, #735361, Sartorius).

As a read-out for cell viability, bulk ATP quantitation with CellTiter-Glo® 2.0 (CTG; #G9243, Promega) to determine the relative number of metabolically active cells per well was performed 72 h after treatment according to the manufactureŕs protocol.

Prior to the drug screen, the suitable cell number to be seeded per well was determined by assessing the proportionality between the number of cells and the luminescence signal for 200, 500, 1,000 and 2,000 cells in the absence of drug treatment. The cell numbers used for the subsequent screening experiment were 250 cells per well for A-673 Dox (–) and 350 cells per well for A-673 Dox (+).

Drug effects were calculated as drug sensitivity scores (DSSasym) using the web-based drug analysis pipeline iTReX^62^.

In analyses of Ribociclib and Palbociclib drug sensitivity screenings of primary patient cultures as determined in the context of the INFORM registry^39^, a culture was considered as responsive if a drug sensitivity score ranged in the highest quartile, whereas samples ranging in the lower 3 quartiles were considered as non-responsive.

### CDK4/6 inhibitor assays in vitro

A-673 and TC-71 (high endogenous 4E-BP1 expression) as well as CHLA-10 and EW-22 (low endogenous 4E-BP1 expression) EwS cells were seeded in a 96-well plate at a density of 0.5 –2× 10^3^ per well. In case of cells containing Dox-inducible constructs, cells were treated with/without Dox (0.5 µg/ml; Sigma-Aldrich). 48h after seeding or pre-incubation with Dox, respectively, CDK4/6 inhibitors (Palbociclib or Ribociclib; Biozol Diagnostica and Hölzel Diagnostika) were added in serially diluted concentrations ranging from 0.0032 to 500 µM. Each well contained an equal concentration of 0.5% DMSO (Sigma-Aldrich). Cells only treated with 0.5% of DMSO served as a control. After 72 h of inhibitor treatment, the plates were assayed on a GloMax® Explorer Multimode Microplate Reader after incubation with Resazurin (16 µg/ml; Sigma-Aldrich) for 4-6 h.

### Mass spectrometry (MS)

Pulse-Chase Labelling using AHA (azidohomoalanine) and SILAC isotope labels for nascent proteome analysis: A-673, SK-N-MC, and TC-71 EwS cells containing either a Dox-inducible non-targeting control shRNA or *EIF4EBP1*-targeting specific shRNAs (sh_4E-BP1_1 or sh_4E-BP1_2) were seeded at a density of 1×10^6^ per 15-cm dish in 20 ml of culture medium and grown with or without Dox (0.5 µg/ml; Sigma-Aldrich) for 96h. Subsequently, cells were washed with pre-warmed PBS and cultivated in 10 ml SILAC RPMI deprivation media (lysine-, arginine-, and methionine-free) (Athenas) for 45 min at 37°C. This ensures the depletion of the intracellular stores of amino acid. The depletion medium was then aspirated and cells were incubated in 13 ml of SILAC RPMI pulse media containing 18.1 mg/l AHA (Jena Bioscience), and either 200 mg/l [13C6, 15N4] L-arginine and 40 mg/l [13C6, 15N2] L-lysine or 200 mg/l [13C6] L-arginine and 40 mg/l [4,4,5,5-D4] L-lysine (Silantes) for 6h at 37°C with 5% CO_2_. Following pulse labelling, cells were washed with PBS and harvested by scraping and centrifugation for 5 min at 400**×** g. Cell pellets were frozen and stored at –80°C until lysis. Enrichment of newly synthesized proteins for translatome analysis and sample preparation for LC-MS/MS measurements: Newly synthesized proteins were enriched using the Click-It alkyne agarose enrichment kit (Thermo Fisher) according to the manufacturer protocol with minor modifications. Cell pellets were lysed in 900 µl of Urea Lysis buffer (200 mM HEPES pH8, 0.5 M NaCl, 4% CHAPS, 8 M Urea), and cell lysates were sonicated on ice using the Benson probe sonicator. Protein concentrations were determined using the BSA kit (Thermo Fisher) and 1 mg of each label pair (+/– Dox) were mixed in a new tube. Volumes were made up to 1,700 µl end volume with 8M Urea lysis buffer. Next, the mixed lysates were combined with 100 µl of the alkyne-agarose slurry and 93 µl of the [3+2] cyclo-addition reaction mixture (10 µl Cu(II)SO4 (200 mM), 62.5 µl tris (hydroxypropyltriazolylmethyl) amine (THPTA, 160 mM), 10 µl aminoguanidine (2 M), 10 µl sodium ascorbate (2 M)), and incubated on a ThermoMixer for 2 h at 40°C. Following the incubation step, the resins were pelleted by centrifugation for 1 min at 2,000**×** *g* and supernatants were discarded. Subsequently, the resins were washed once with 2 ml of MQ water and pelleted by another centrifugation step. Next, the resins were resuspended in 2 ml of the 1-step reduction/alkylation mixture (10 mM Tris(2-carboxyethyl)phosphine, 40 mM 2-Chloroacetamide in SDS wash buffer) and incubated on a ThermoMixer for 15 min at 70 °C and another 15 min at 20 °C. Following the reduction/alkylation step, the resins were transferred into spin columns (BioRad) and placed on a P10 tip-box for the subsequent steps. The resins were consecutively washed five times with 1 ml SDS-Wash buffer (100 mM Tris-HCl pH 8.0, 1% SDS, 250 mM NaCl, 5 mM EDTA), once with MQ water, five times with Guanidine-Wash buffer (100 mM Tris-HCl pH 8.0, 6M Guanidine-HCl), and 5**×** with Acetonitrile-Wash buffer (20% Acetonitrile, ULCMS in Water ULCMS). Subsequently, the resins were resuspended in 200 µl Digestion buffer (100 mM Tris-HCl pH 8, 2 mM CaCl2, 5% Acetonitrile), transferred to new tubes, and subjected to proteolytic digestion by adding 1 µg of trypsin on a ThermoMixer for 16h at 37 °C. Following the digestion step, the resins were pelleted by centrifugation for 5 min at 2,000**×** g, the supernatants containing the digested peptides were transferred to new tubes and acidified with formic acid (FA) to 1% end concentration. For the peptide clean-up step, Oasis PRiME HLB µElution Plates (Waters) were used according to the manufacturer protocol. Briefly, the digested peptides were transferred to the Oasis plate and washed consecutively with 750 µl, 250 µl, and 100 µl of 1% FA. Peptides were eluted in a 96-well plate by adding 70 µl of elution solution (60% MeOH, 1% FA, 39% Water ULCMS) and another 50 µl of 100% MeOH. The eluted peptides were then dried down using a SpeedVac, resuspended in 0.1% TFA, and samples were subjected to HPLC fractionation and mass spectrometry analysis. LC-MS/MS analysis: Samples were analyzed on an Orbitrap Fusion mass spectrometer (Thermo Fisher) coupled with an Easy-nLC 1200 system (Thermo Fisher). Chromatographic separation was carried out using an Acclaim Pepmap RSLC trap column (100 µm **×** 2 cm, 5 µm particles, 100 Å pores, C18, Thermo Fisher), and nanoEase M/Z Peptide BEH analytical column, (75 µm **×** 250 mm, 1.7 µm particles, 130Å pores, C18, Waters). Peptides elution gradient was set to 105 min at flow rate of 300 nl/min using solvent A (0.1% formic acid in ULCM grade water) and solvent B (0.1% formic acid in 80% acetonitrile and 19.9% ULCM grade water). Samples were injected into the mass spectrometer using a 10 µm Picotip coated fused silica emitter (New Objective). The Orbitrap-Fusion mass spectrometer was operating in positive mode. Acquisition was carried out in data-dependent acquisition (DDA) mode. The MS1 scan was detected in orbitrap mode at 60,000 FWHM resolution, AGC target 1E6, scan range (m/z) was 375–1,500 DA, and maximal injection time was set to 50 ms. The intensity threshold HCD-fragmentation was set to 5E3. MS2 detection was acquired in centroid mode with an HCD collision energy of 33% in an ion trap detector, an isolation window (m/z) of 1.6 Da, AGC target 1E4, and maximal injection time of 50 ms. Data analysis: The raw files were analyzed using MaxQuant (version 1.6.10.43). The MS/MS spectra were searched through the integrated Andromeda search engine against Homo Sapiens UniProt proteome database. Multiplicity of labels was set to 3. Arg6/Lys4 and Arg10/Lys8 were selected as medium and heavy labels, respectively. Cysteine carbamidomethylation was selected for fixed modification. Methionine oxidation, protein N-terminal acetylation, replacement of methionine by AHA and conversion of AHA to homoserine (HS) and diaminobutyrate (DAB) were selected for variable modifications. Re-quantify and match between runs (match time window: 0.4 min) functions were enabled. For the digestion, Trypsin/P was selected, and the maximal number of mis-cleavages was set to 2. The minimal peptide length was set to 7 amino acids and FDR for peptide and protein identification was set to 0.01. For protein identification, at least one unique peptide was required. Minimum ratio count for label-based protein quantification was set to 2. Normalized H/M ratios derived from the ProteinGroups file were further processed and subjected to statistical analysis. FDR was set to >0.05 and S0 to 0.1. MS profiling was performed in biological triplicates for every cell line/construct/condition. This process identified 9,508 proteins through MS analysis, of which 4,335 common proteins across all cell lines and constructs with at least one value per replicate group were used for downstream analyses.

For analysis of differential protein expression, log_2_ fold changes (log_2_FCs) between conditions with and without Dox treatment were calculated for every cell line, every construct, and every biological replicate. For every cell line/construct log2FCs were normalized to its respective shCtr. Next, a global median log_2_FC across all biological replicates was calculated for every protein. Proteins were only considered as being directly regulated by 4E-BP1 when their mRNA |log_2_ FCs| were <0.5.

### Xenotransplantation experiments

To assess local tumor growth of subcutaneous xenografts *in vivo*, 2.5×10^6^ A-673 or TC-71 EwS cells containing either a Dox-inducible negative control shRNA or specific shRNAs against *EIF4EBP1* were injected subcutaneously with a 1:1 mix of PBS (Biochrom) and Geltrex (LDEV-Free Reduced Growth Factor Basement Membrane Matrix, Thermo Fisher Scientific; max volume 100 µl) in the right flanks of 4–8 week old NSG mice following routine protocols^63^. When tumors were first palpable, mice were randomized to the control group (17.5 mg/ml sucrose (Sigma-Aldrich) in drinking water) or the treatment group (2 mg/ml Dox Beladox, bela-pharm) and 50 mg/ml sucrose (Sigma-Aldrich) in drinking water. According to the average water intake of mice in the different treatment groups, the concentration of sucrose in the different treatment groups has been adapted: due to the bitter taste of Dox, the mice that receive Dox via their drinking water in average have a lower water intake, making a higher concentration of sucrose necessary to ensure equal intake of sucrose per mouse across the different treatment groups. Tumor size was measured with a calliper every two days and tumor volume was calculated as *V=a×b^2^/2* with *a* being the largest diameter and *b* the smallest. Right before tumors reached the maximum size of 15 mm in one dimension (event), the respective mice were sacrificed by cervical dislocation. Other specific humane endpoints were determined as follows: invasive tumor growth leading to functional impairment or pain, ulcerating tumor (or fluid externalization), prolonged obstipation, abdominal distention, peritonitis, ascites, bloody diarrhea, palpable abdominal tumor mass with additional signs of pain, total relief, or medium paresis of a limb. General humane endpoints were determined as follows: Loss of 20% body weight, apathy, piloerection, self-isolation, aggressivity, automutilation, unphysiological or reduced movements/positioning, abnormal breathing, and reaching of a maximum observation period of 12 months.

For analysis of EwS xenograft growth in bone, TC-71/TR/sh4E-BP1_2 EwS cells were orthotopically injected into the proximal tibial plateau of NSG mice. One day before injection, mice were pre-treated with 800 mg/kg mouse weight/d Metamizole in drinking water as analgesia. On the day of injection, mice were anesthetized with inhaled isoflurane (2.5% in volume) and their eyes were protected with Bepanthen eye cream. After disinfection of the injection site, 2×10^5^ cells/20 µl were directly injected with a 30 G needle (Hamilton) into the right proximal tibia. For pain prophylaxis after intraosseous injection, mice were treated with Metamizole in drinking water (800 mg/kg mouse weight/d). The first day after injection of tumor cells, mice were randomized in two groups of which one received henceforth 2 mg/ml Dox (BelaDox, Bela-pharm) dissolved in drinking water containing 5% sucrose (Sigma-Aldrich) to induce sh4E-BP1_2 expression, whereas the other group only received 5% sucrose. All mice were closely monitored routinely every two days and tumor growth was evaluated with a caliper. All tumor-bearing mice were sacrificed by cervical dislocation when the mice exhibited first signs of limping at the injected leg (event) or reached any humane endpoint as listed above.

For CDK4/6 inhibitor treatment *in vivo*, A-673/TR/sh4E-BP1_1 cells were injected subcutaneously as described above. As soon as the tumors were palpable, mice were subjected to either the vehicle (DSMO) or the treatment group (Palbociclib, LC Laboratories, 100 mg/kg), whereby each group was treated with or without addition of Dox to the drinking water (Beladox, bela-pharm, 2 mg/ml). Palbociclib was administered by oral gavage for 28 days, with a break of 2 days every 5 days of treatment. The experimental endpoint was predetermined as 28 days after first injection of either inhibitor, or if humane endpoints as described above were reached before. To examine the number of mitoses within the tumors, hematoxylin and eosin (HE) stained slides of the respective tumors were examined and the number of mitoses were quantified as described below in section ‘Histology’. Animal experiments were approved by the government of Upper Bavaria and North Baden and conducted in accordance with ARRIVE guidelines, recommendations of the European Community (86/609/EEC), and United Kingdom Coordinating Committee on Cancer Research (UKCCCR) guidelines for the welfare and use of animals in cancer research.

### Methylation arrays and CNV analysis

A minimum of 500 ng of high-quality DNA from 100 EwS FFPE samples was used for methylation and CNV analysis with the Infinium Human Methylation 450K BeadChip (EPIC array, Illumina). This method can analyze 864,928 CpGs including main CpG islands and CpG sites outside of CpG islands. Raw methylation and CNV data were essentially processed as previously reported^64^. Based on the methylome, we used a previously described sarcoma classifier^64^, which assigns a score to each sample indicating the similarity of the respective methylome with methylation patterns of samples from known entities. CNVs were also assessed by this method and analyzed at both, chromosomal and specific locus level. Scores for specific positions were visualized in the Integrative Genomic Viewer (IGV).

For analysis regarding the association between chr8 gain and *EIF4EBP1* expression (as assessed by RNA-seq), only samples were analyzed which were clearly possible to separate into either a group showing global chr8 gain or no chr8 gain. Samples with partial gain were not considered.

For determining chr8 CNA status in tumor entities other than EwS we analyzed TCGA SNP array 6.0 pre-segmented data using TCGAbiolinks R package^65^ (Masked Copy Number segment, GRCh38.p0). For downstream analysis we selected all primary tumor samples for which clinical annotation was complete and SNP array and RNA-seq datasets existed, we applied re-segmentation algorithm by CNApp^66^ to classify samples as chr8 gain or no gain. Classification criteria for whole chr8 gain were log2 segment mean > 0.2 and the longest segment spanning ≥ 75% of total chr8 length (chr8q length 68%). To compare both groups (chr8 gain vs. no gain) regarding overall patient survival we performed a Kaplan Meier analysis using the survival R package^67,68^. Significance levels were calculated using the log-rank test.

### RNA sequencing (RNA-seq)

RNA quality was assessed by the Agilent 2100 Bioanalyzer (Agilent Technologies) before library preparation. 2 µg of estimated high-quality RNA was prepared for sequencing. Libraries were prepared according to the TruSeq RNA Exome (Illumina) protocol. Library preparation workflow included purification and fragmentation of mRNA, first and second strand cDNA synthesis, end repair, 3’ends adenylation, adapters ligation, PCR amplification, library quantification, normalization and libraries pooling. Cleanup cycles are introduced between the mentioned steps. Sequencing was performed on a NextSeq 550 / NovaSeq 600 Sequencer (Illumina). NextSeq 550 was performed with 75 bp ‘paired-end’ sequencing technology employing high-output flow cells. Calculations of RNA counts were performed as previously described^69,70^.

### Gene expression microarrays

A-673, TC-71, and SK-N-MC EwS cells containing either a Dox-inducible non-targeting control shRNA (shCtr) or *EIF4EBP1*-targeting specific shRNAs (sh4E-BP1_1 or sh4E-BP1_2) were seeded in T25 flasks (TPP) and treated either with or without Dox (0.5 µg/ml; Sigma-Aldrich) for 96 h. Thereafter, total RNA was extracted from one biological replicate for each condition with the Nucleospin II kit from Macherey-Nagel and transcriptome profiled at IMGM laboratories (Martinsried, Germany). RNA quality was assessed with a Bioanalyzer and samples with RNA integrity numbers (RIN) > 9 were hybridized to Human Affymetrix Clariom D microarrays. Data were quantile normalized and summarized with Transcriptome Analysis Console (v4.0; Thermo Fisher Scientific) using the SST-RMA algorithm. Annotation of the data was performed using the Affymetrix library for Clariom D Array (version 2, human) at gene level. Differentially expressed genes across shRNAs and cell lines were identified as follows: First, normalized gene expression signal was log_2_ transformed. To avoid false discovery artifacts due to the detection of only minimally expressed genes, we excluded all genes with a lower or just minimally higher gene expression signal than that observed for *ERG*, which is known to be virtually not expressed in *EWSR1::FLI1* positive EwS cell lines^18^. A log_2_FC was calculated for every cell line/construct and normalized to its respective shCtr. Normalized log_2_FCs were summarized across cell lines/construct as median log_2_FC.

### Analysis of publicly available gene expression data and patient survival analysis

Microarray data of 196 EwS tumors (Cohort 1) (GSE63157, GSE34620, GSE12102, GSE17618 and unpublished data^71–74^) for which well-curated clinical annotations were available were downloaded from the Gene Expression Omnibus (GEO). The data were either generated on Affymetrix HG-U133Plus2.0 or on Affymetrix HuEx-1.0-st microarray chips and were normalized separately by RMA using custom brainarray chip description files (CDF, v20) as previously described^75^. Batch effects were removed using ComBat^76^. Tumor purity was calculated using ESTIMATE^77^ and only samples with a tumor purity >60% corresponding to The Cancer Genome Atlas (TCGA) standard (http://cancergenome.nih.gov/cancersselected/biospeccriteria) were kept for further analyses.

Samples were stratified by their quartile intra-tumoral gene expression levels. Significance levels were calculated with a Mantel-Haenszel test (calculated using GraphPad Prism version 9). *P*-values <0.05 were considered as statistically significant.

For analyzing the effect of *EIF4EBP1* expression level on clinical outcome in tumor entities other than EwS we analyzed TCGA RNA-seq unnormalized counts data using TCGAbiolinks R package^65^ (STAR - Counts, GRCh38.p0). For downstream analysis we selected all primary tumor samples for which clinical annotation was complete and SNP array and RNA-seq datasets existed. We kept all genes with at least 50% samples showing counts greater than 0. Using DESeq2^78^ for downstream analysis we applied variance stabilizing transformation algorithm on the counts matrix to produce normalized counts on the log2 scale. To investigate the association between *EIF4EBP1* expression and overall patient survival we performed Kaplan Meier analyses using the best percentile approach^67^. To determine the optimal threshold based on *EIF4EBP1* expression as a continuous variable (vst-counts), we used maximally selected rank statistics from the maxstat R package^79^. Significance levels were calculated using the log-rank test.

### Fast gene-set enrichment analysis (fGSEA) and single-sample GSEA (ssGSEA)

fGSEA was performed using the FGSEA R package (v 4.1.3) based on Gene Ontology (GO) biological processes terms from MSigDB (c5.all.v7.5.1 symbols.gmt, c2.all.v7.5.1 symbols.gmt, c2.cgp.v7.5.1 symbols.gmt) and GO terms were filtered for statistical significance (adjusted *P*<0.05)^80,81^.

Using the Affymetrix gene expression dataset comprising 196 EwS patients (Cohort 1), enrichment of gene sets that are among *EIF4EBP1* co-regulated genes were identified by ranking of Pearsońs correlation coefficient of the expression of each gene with *EIF4EBP1* expression and performance of a pre-ranked fGSEA.

Using the RNA-seq gene expression dataset comprising 100 primary EwS patients (Cohort 2), enrichment of gene sets that are among *EIF4EBP1* co-regulated genes and among chr8 regulated genes were identified by performing a pre-ranked fGSEA on a) ranked Pearsońs correlation coefficient between the expression of each gene with *EIF4EBP1* expression and on b) ranked mean log_2_FC as calculated for each gene between EwS with and without chr8 gain.

Employing an integrated dataset inferred from MS protein expression data and microarray gene expression data of A-673, SK-N-MC, and TC-71 EwS cell lines with/without 4E-BP1 knockdown containing differentially expressed proteins upon 4E-BP1 knockdown, all proteins were ranked by their median log_2_ FC and a pre-ranked fGSEA was performed.

For inferring chr8 gene expression enrichment from gene expression data we applied single sample GSEA (ssGSEA)^82,83^ on a EwS patient cohort of 196 tumor samples (Cohort 1) using gene sets from Ensembl 103 genes in cytogenetic band chr8 (Molecular Signatures Database (v7.4 MSigDB/c1.all.v7.5.symbols.gmt)). Genes considered for chr8 gene expression signature are listed in **Supplementary Table 11**. Based on the enrichment levels (ssGSEA scores) of chr8, we performed hierarchical clustering (pheatmap R package, version 1.0.12, https://cran.r-project.org/web/packages/pheatmap/pheatmap.pdf) to divide the EwS samples into a chr8 high and low gene expression group. To refine the chr8 enrichment approach by excluding samples with other recurrent CNVs, the same approach as for chr8 was applied with gene sets reflecting chr1q, chr12, and chr16. Samples were assigned as only chr8 high when none of the other recurrent CNVs were present, and as only chr8 low when none recurrent CNVs were present. This resulted in 117 EwS patients (Cohort 1, chr8 focus) which were included in this refined analysis. To compare both groups (chr8 gene expression high vs. low) regarding overall patient survival we performed a Kaplan Meier analysis. Significance levels were calculated with Mantel-Haenszel test (GraphPad Prism version 9).

### Position Related Data Analysis (PREDA)

To validate appropriate clustering of Cohort 1 tumor samples into chr8 high and low gene expression groups we determined differentially expressed genes (DEGs) between both clusters using Limma^84^ and Position Related Data Analysis (PREDA)^85^ on iDEP^86^ platform to map respective DEGs (FDR cut-off < 0.01) onto chromosomes. Using the Limma package^84^, also DEGs of the mTOR signaling pathway were depicted.

### Cancer Dependency Map (DepMap) analyses

To identify potential therapeutic targets for EwS cells, we leveraged existing curated cancer Dependencey Map (DepMap, Broad Institute) data^87–89^. For each gene, moderated estimates of the differences between the means of gene dependency effects across EwS (*n*=23) versus all other cell lines (*n*=1077) were plotted against its statistical significance. Q-values were calculated to adjust for multiple testing.

### Histology

HE-staining of EwS xenografts was performed according to routine protocols. Mitoses of EwS xenografts were quantified in HE-stained slides by two blinded observers in 5 high-power fields per sample. Mitoses per sample were determined by mean of total 10 counted high-power fields across both observers.

### Statistics and software

Statistical data analysis was performed using GraphPad PRISM 9 (GraphPad Software Inc., CA, USA) or with R (version 4.2.0) on the raw data. If not otherwise specified in the figure legends, comparison of two groups in functional *in vitro* experiments was carried out using a two-tailed Mann-Whitney test. If not otherwise specified in the figure legends, data are presented as dot plots with horizontal bars representing means, and whiskers representing the standard error of the mean (SEM). Sample size for all *in vitro* experiments was chosen empirically with at least 3 biological replicates. In Kaplan-Meier overall survival analyses, curves were calculated from all individual survival times of patients. Curves were compared by Mantel-Haenszel test to detect significant differences between the groups. For batch analyses of patient survival, the in-house custom code software GenEx was used, using the Mantel-Haenszel test for *P*-value calculation. Pearsońs correlation coefficients were calculated using Microsoft Excel or GraphPad PRISM 9 (GraphPad Software Inc., CA, USA). For *in vivo* experiments, sample size was predetermined using power calculations with β = 0.8 and α = 0.05 based on preliminary data and in compliance with the 3R system (replacement, reduction, refinement). Kaplan-Meier analyses of event-free survival (*in vivo* experiments) were carried out using GraphPad PRISM 9 (GraphPad Software Inc.). The definition of the type of events is given in the corresponding figure legends, but generally corresponds to humane experimental endpoints as defined above.

## ACKNOWLEGEMENTS

We thank Dr. Aruna Marchetto for her help in western blot assays. We thank Dr. Clemens M. Lechner for statistical advice. Support by the DKFZ Light Microscopy Facility is gratefully acknowledged.

## AUTHOR CONTRIBUTIONS

C.M.F. and A.C.E. performed functional *in vitro* and *in vivo* experiments, as well as bioinformatic and histological analyses, analyzed and interpreted all data, designed all figures, and wrote the paper. M.F.O. performed functional experiments, helped in shRNA design and lentiviral transduction of cell lines. K.A. performed MS and analyzed MS data. J.S. and A.Y. helped in dataset curation. J.L. helped in conduction of *in vivo* experiments. T.L.B.H., M.M.L.K., E.V., A.K.C. assisted in experimental procedures. R.W. performed cloning of vectors. F.W. performed analyses of histology. F.Z. assisted in experimental procedures. S.O., R.I., A.B. helped in performance of *in vivo* experiments. J.A., A.S., W.H., U.D. provided clinical information. M.S. provided financial support and laboratory infrastructure. O.W., I.O., H.P., and A.L. performed drug screens. S.M.P., G.L. provided biological and technical guidance. L.R.-P. conducted methylation arrays and performed bioinformatic analyses. J.K. provided financial support and laboratory infrastructure for performance of MS. F.-C.A. carried out functional *in vitro* and *in vivo* experiments. J.M. coordinated and supervised the study, provided biological and technical guidance, performed functional experiments as well as bioinformatic and histological analyses, analyzed and interpreted all data, wrote the paper, and provided financial support. T.G.P.G designed, coordinated, and supervised the study, provided biological and technical guidance, analyzed and interpreted all data, wrote the paper and provided financial support and laboratory infrastructure. All authors read and approved the final manuscript.

---

**Supplementary Figure 1:**
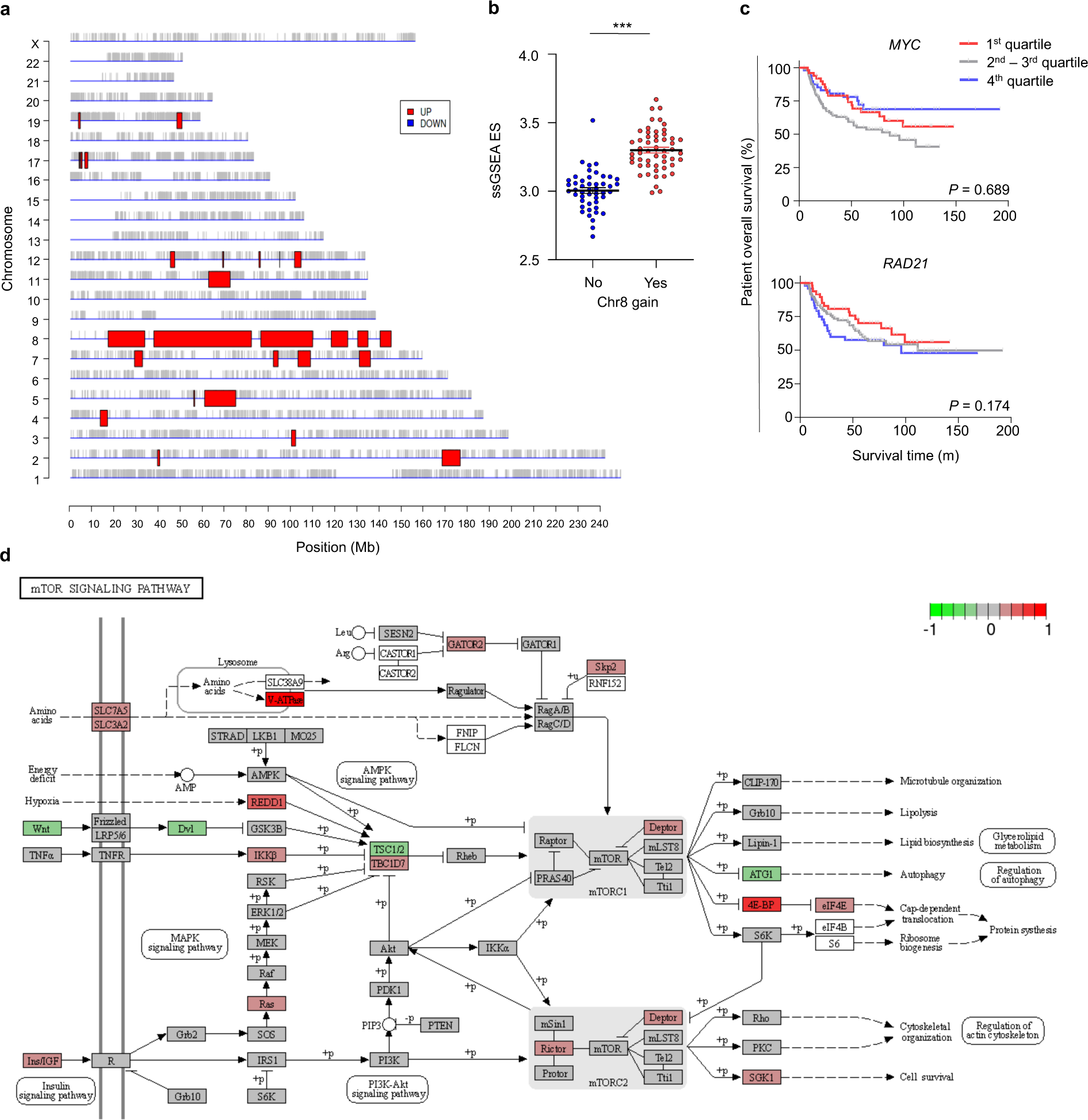
Model validation, chromosomal location, and mTOR pathway representation of genes differentially upregulated in the chr8 high gene expression signature group in EwS. **a** Differentially expressed genes (DEGs) between the chr8 high and low gene enrichment clusters mapped onto chromosome positions using Position Related Data Analysis (PREDA). **b** ssGSEA enrichment scores for chr8 gene expression enrichment as measured by RNA-seq in 100 primary EwS (Cohort 2) depending on the presence of factual chr8 gain as determined by methylation array. **c** Kaplan-Meier overall survival analysis of 196 EwS patients (Cohort 1) stratified by quartile *MYC* or RAD21 expression. *P-*values determined by Mantel-Haenszel test. **d** DEGs between the chr8 high and low gene expression cluster in Cohort 1 within the mTOR signaling pathway. \*\*\**P* < 0.001, *P*-values determined via two-tailed Mann-Whitney test.

**Supplementary Figure 2:**
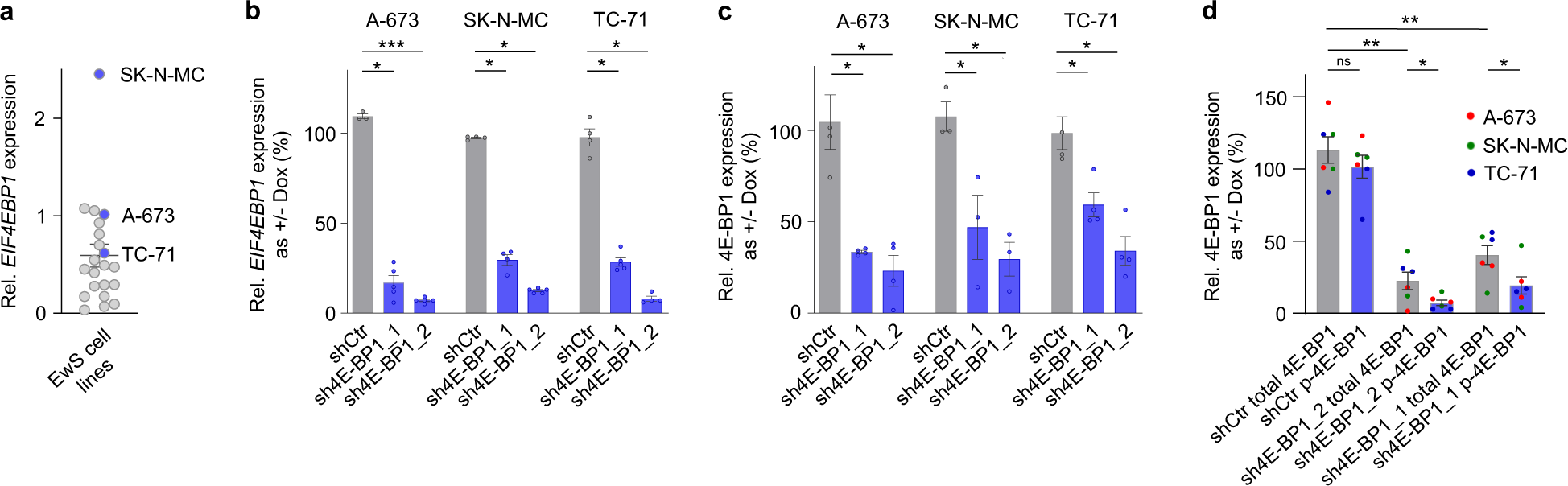
4E-BP1 drives a multifunctional proliferation-associated proteomic network. **a** Relative *EIF4EBP1* expression in 21 wildtype EwS cell lines as determined by qRT-PCR. *EIF4EBP1* expression of each cell line is normalized to that of A-673. **b** Relative *EIF4EBP1* expression as assessed by qRT-PCR in A-673, SK-N-MC, and TC-71 cells containing either Dox-inducible specific shRNA constructs directed against *EIF4EBP1* (sh4E-BP1_1 or sh4E-BP1_2) or a non-targeting shControl (shCtr). Cells were grown either with or without Dox for 96 h. Horizontal bars represent means, and whiskers represent the SEM, n≥3 biologically independent experiments. **c** Relative 4E-BP1 expression as assessed by quantified western blotting in A-673, SK-N-MC, and TC-71 cells containing either Dox-inducible specific shRNA constructs directed against *EIF4EBP1* (sh4E-BP1_1 or sh4E-BP1_2) or a non-targeting shControl (shCtr). Cells were grown either with or without Dox for 96 h. *P-*values determined via one-tailed Mantel-Haenszel test. **d** Relative total and phospho (Ser65) 4E-BP1 expression as assessed by quantified western blotting in A-673, SK-N-MC, and TC-71 cells containing either Dox-inducible specific shRNA constructs directed against *EIF4EBP1* (sh4E-BP1_1 or sh4E-BP1_2) or a non-targeting shControl (shCtr). Cells were grown either with or without Dox for 96 h. *P-* values determined via one-tailed Mantel-Haenszel test.

**Supplementary Figure 3:**
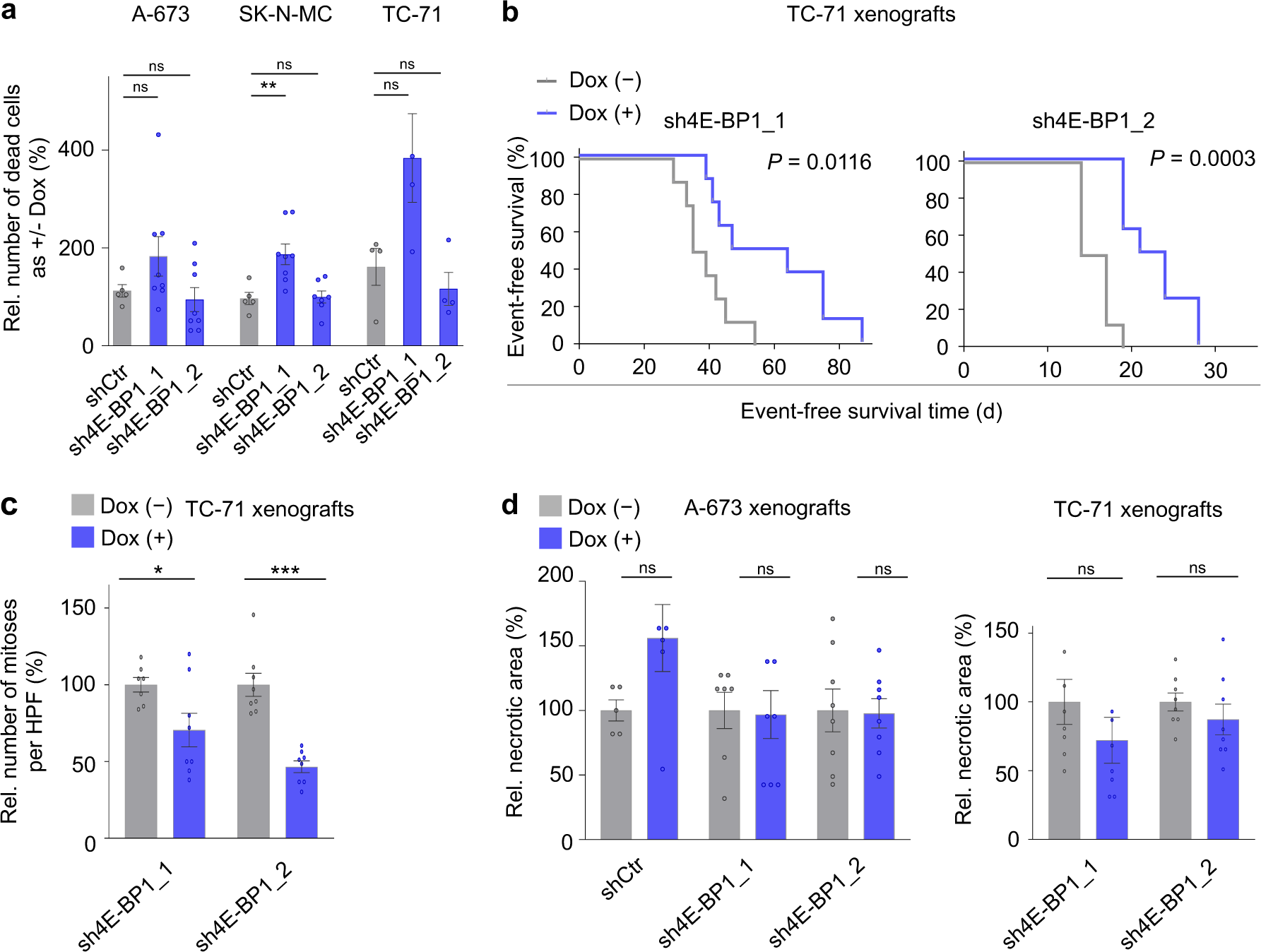
RNAi-mediated knockdown of 4E-BP1 inhibits EwS growth. **a** Relative number of dead cells as assessed by Trypan blue exclusion in A-673, SK-N-MC, and TC-71 cells containing either Dox-inducible specific shRNA constructs directed against *EIF4EBP1* (sh4E-BP1_1 or sh4E-BP1_2) or a non-targeting shControl (shCtr). Cells were grown either with or without Dox for 96 h. Horizontal bars represent means, and whiskers represent the SEM, n≥4 biologically independent experiments. **b** Kaplan-Meier analysis of event-free survival of NSG mice xenografted with TC-71 cells containing Dox-inducible specific shRNA constructs directed against *EIF4EBP1* (sh4E-BP1_1 or sh4E-BP1_2). Once tumors were palpable, mice were randomized and treated with either vehicle (–) or Dox (+), n =8 animals per condition. An ‘event’ was recorded when tumors reached a size maximum of 15 mm in one dimension. *P-*values determined via Mantel-Haenszel test. **c** Quantification of mitoses in HE-stained slides of xenografts described in (b). Five high-power fields (HPF) were counted per sample. Horizontal bars represent means, and whiskers represent the SEM, n ≥ 7 samples per condition. **d** Quantification of necrotic area on HE-stained slides of A-673 and TC-71 xenografts described in (Fig. 3d, Suppl. Fig. 3b). Five high-power fields (HPF) were analyzed per sample. Horizontal bars represent means, and whiskers represent the SEM, n≥5 samples per condition.

**Supplementary Figure 4:**
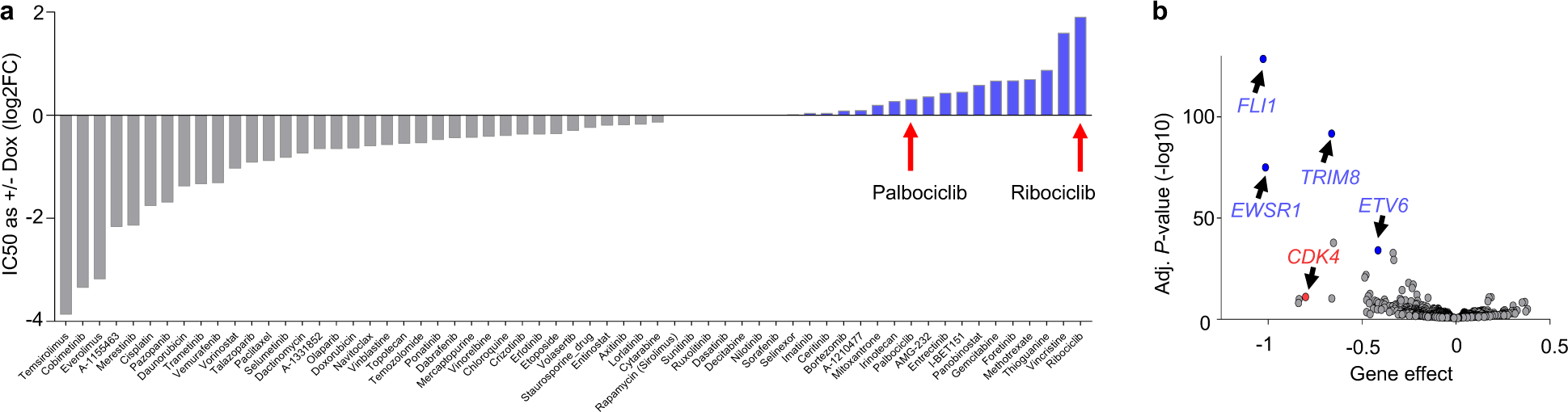
High 4E-BP1 expression sensitizes for targeted CDK4/6 inhibitor treatment with Palbociclib and Ribociclib. **a** IC50 analysis of 3D culture drug screening data of A-673 EwS cells containing a Dox-inducible shRNA directed against 4E-BP1 and treated with/without Dox and respective indicated inhibitors in serially increasing concentrations. **b** Volcano plot showing gene dependency effects of indicated genes in EwS cell lines as compared to all non-EwS cell lines with respective individual statistical significance values (-log10 adj. *P*-value).

